# Rapid multicomponent bioluminescence imaging via substrate unmixing

**DOI:** 10.1101/811026

**Authors:** Colin M. Rathbun, Anastasia A. Ionkina, Zi Yao, Krysten A. Jones, William B. Porterfield, Jennifer A. Prescher

## Abstract

Engineered luciferases and luciferins have dramatically expanded the scope of bioluminescence imaging in recent years. Multicomponent tracking remains challenging, though, due to a lack of streamlined methods to visualize combinations of bioluminescent reporters. Here we report a strategy for rapid, multiplexed imaging with a wide range of luciferases and luciferins. Sequential addition of orthogonal luciferins, followed by substrate unmixing, enabled facile detection of multiple luciferases *in vitro* and *in vivo*. Multicomponent imaging in mice was also achieved on the minutes-to-hours time scale, a vast improvement over conventional protocols.

## MAIN

Bioluminescence imaging (BLI) is a popular technique for tracing cells and other biological features in heterogeneous environments, including whole animals.^1^ BLI relies on genetically encoded luciferase enzymes and luciferin substrates for photon production. Because mammalian tissues do not normally glow, BLI enables sensitive imaging *in vivo*. As few as 1-10 cells can be reliably detected using optimized probes in subcutaneous models. For these reasons, BLI has long been a go-to technique for monitoring physiological processes in rodents.^2-4^ More recent advances are further enabling studies in non-human primates and other large organisms.^5-6^

While ubiquitous, BLI applications *in vivo* typically track only one cell type or feature at a time.^2^ A spectrum of bioluminescent reporters exists, but discriminating them based on wavelength alone is challenging.^3^ In fact, most applications featuring two BLI reporters combine firefly luciferase (Fluc) and *Renilla* luciferase (Rluc), two enzymes that use different substrates.^7-8^ Multicomponent imaging with Fluc and Rluc *in vivo*, though, often requires multiple hours, if not days. The bioluminescent substrates are typically administered at saturating doses to maximize photon output, and the first probe must clear prior to administering the second. In principle, dozens of other naturally occurring luciferases and luciferins could be employed for multiplexed BLI.^9^ In practice, though, most of these enzymes and substrates are ill-suited for *in vivo* use.

We aimed to address the need for better probes and more practical imaging protocols for multicomponent BLI. Our approach builds on the expanding toolbox of orthogonal bioluminescent reagents.^3^ These probes comprise genetically modified luciferases that are responsive to chemically unique luciferins.^5,10-13^ A handful of the orthogonal enzymes can be readily discriminated in cells and mouse models based on selective substrate use.^10-11^ Like Fluc and Rluc, though, traditional applications of the engineered reporters require long times (hours-to-days) for complete image capture.^11^ Here we report a method to rapidly resolve orthogonal bioluminescent probes on the minutes-to-hours time scale. The approach is broadly applicable and scalable to multiple reporters, enabling facile multiplexed bioluminescence imaging.

We hypothesized that rapid BLI could be achieved via sequential substrate administration and serial image acquisition. Light outputs would build over time, with each luciferin application resulting in stronger signal. A final processing step^14^ would unmix the images (Fig. 1a). This approach fundamentally differs from conventional optical imaging, in that wavelength, lifetime, and other traditional parameters are less significant. The probes must simply be substrate resolved (i.e., orthogonal) and intensity resolved (Fig 1b). Substrate resolution minimizes cross talk between the probes. Intensity resolution ensures that signal can be “layered in”: as successively brighter probes are imaged, residual signal from dimmer, earlier images becomes part of the background. No time is required for substrate clearance. If the bioluminescent probes are not intensity resolved, the targets are indistinguishable. Similar “layering” approaches have vastly expanded fluorescence detection of gene transcripts^15^ and other cellular features^16-17^ in recent years.

**Figure 1.**
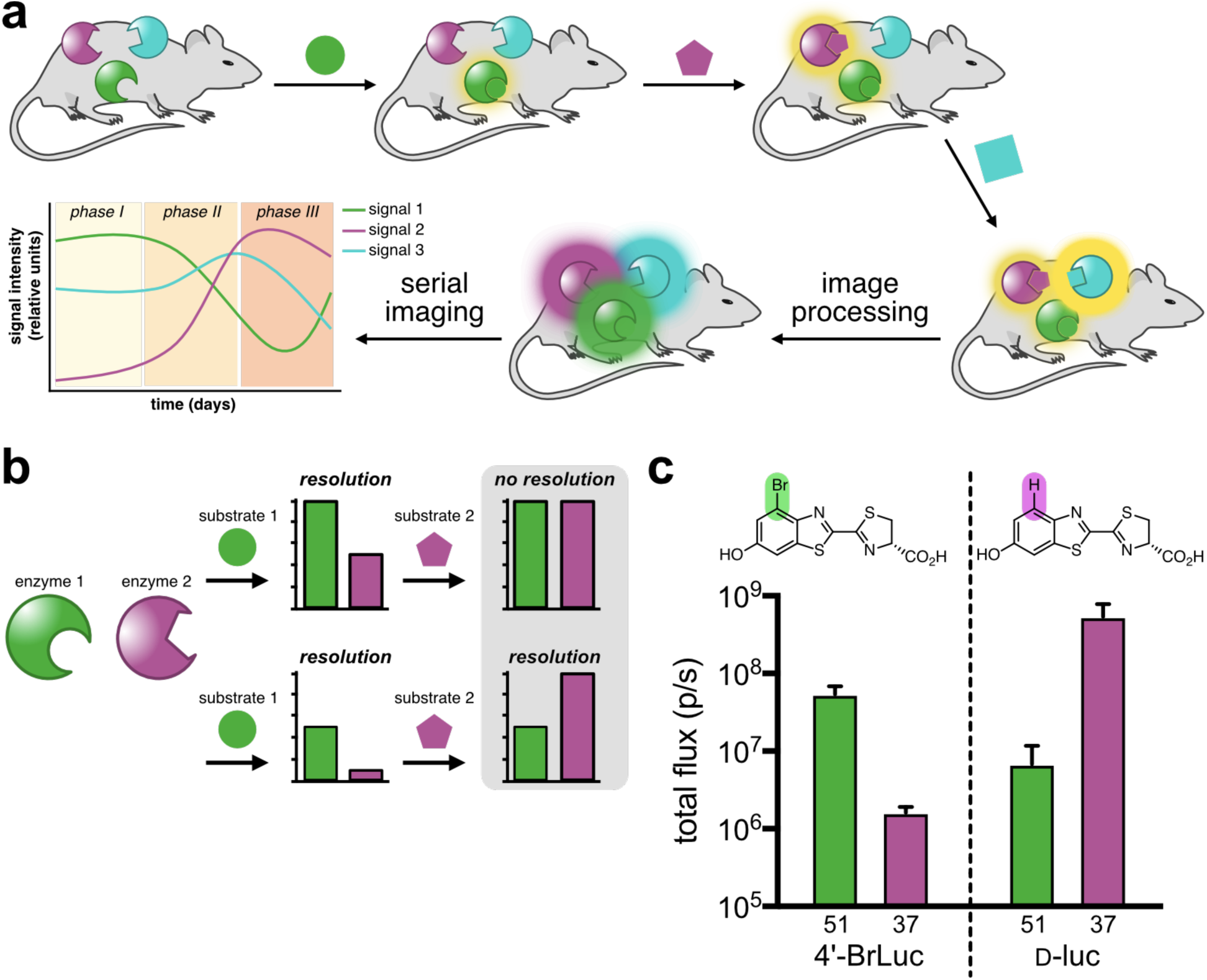
Multicomponent bioluminescence imaging via serial substrate addition and unmixing. (a) Sequential application of orthogonal luciferins (shapes) to illuminate multiple luciferase reporters *in vivo*. Linear unmixing algorithms can deconvolute substrate signatures, enabling rapid and dynamic readouts of biological processes. (b) Bioluminescent probes must be orthogonal and exhibit differential emission intensities for successful unmixing. When both probes are equally “bright” (top), no resolution is possible. Probes of varying intensity (bottom) can be distinguished when the dimmest probe is administered first. (c) An orthogonal bioluminescent pair. Mutants 51 and 37 can be resolved using 4’-BrLuc (100 µM) or D-luc (100 µM). Error bars represent the standard error of the mean for *n* = 3 experiments.

To identify suitable probes for rapid BLI, we focused on two previously reported orthogonal luciferins: 4’-BrLuc and D-luciferin (D-luc, Fig. 1c).^11^ These substrates are bright, bioavailable, and accessible in large quantities. We also previously identified mutant luciferases that could differentiate the analogs.^11^ While orthogonal, these pairs were engineered for maximum brightness and not built with intensity resolution in mind. To identify orthogonal luciferases that exhibited a *range* of photon outputs, we screened a small panel of mutants known to process C4’-modified analogs (Fig S1). Screens were performed both *in vitro* and *in vivo*. In the latter case, DB7 cells stably expressing mutant luciferases were implanted in mice. 4’-BrLuc and D-luc were administered sequentially. The most orthogonal and intensity resolved pair comprised mutants 51 and 37 (Fig S2), which we subsequently named Pecan and Cashew, respectively. Bioluminescent signal from 4’-BrLuc/Pecan is orders of magnitude lower than D-luc/Cashew, meaning that the two pairs are intensity resolved and amenable to rapid sequential imaging (Figs 1c and S2). 4’-BrLuc can be administered first (to illuminate Pecan), followed immediately by D-luc (to illuminate Cashew). Since Cashew signal is brighter than Pecan, the images can be readily unmixed.

To evaluate the reporters for rapid multicomponent BLI, we performed a series of *in vitro* experiments. Pecan and Cashew-expressing cells were ruptured, and lysates were distributed across black-well plates. 4’-BrLuc was initially administered to each well, and an image of the plate was acquired (Fig 2a). D-Luc was then immediately added to the same wells, and a second image was acquired. Because light output from Cashew is brighter, the Pecan signal falls entirely within the noise of the second image. False colors were assigned to each reporter. The images were then overlaid and a linear unmixing algorithm was employed to determine the relative quantities of each mutant. In all cases, the measured signal correlated linearly with probe concentration (Fig 2b). Signal outputs from mixed lysate samples were also co-linear with samples comprising a single luciferase, indicating minimal signal crosstalk (as expected for orthogonal probes). Rapid imaging required the dimmest reporter to be visualized first (Fig S3). The unmixing algorithm is not necessary when the second target is more abundant than the first.^15^ When the second target (associated with the “brightest” luciferase) is in low abundance, though, the algorithm is crucial for proper image interpretation. Unmixing ensures that residual signal from the first image is eliminated and doesn’t interfere with the subsequent image (Fig S4 and Supplementary Discussion). Since the relative abundance of multiple targets is unknown in a given experiment, the algorithm would be employed in all cases by the end user. The “layering in” approach was applicable to several classes of luciferases and luciferins *in vitro* (Fig S5). Multiple mammalian cell populations could also be visualized in a single imaging session (Fig S6).

**Figure 2.**
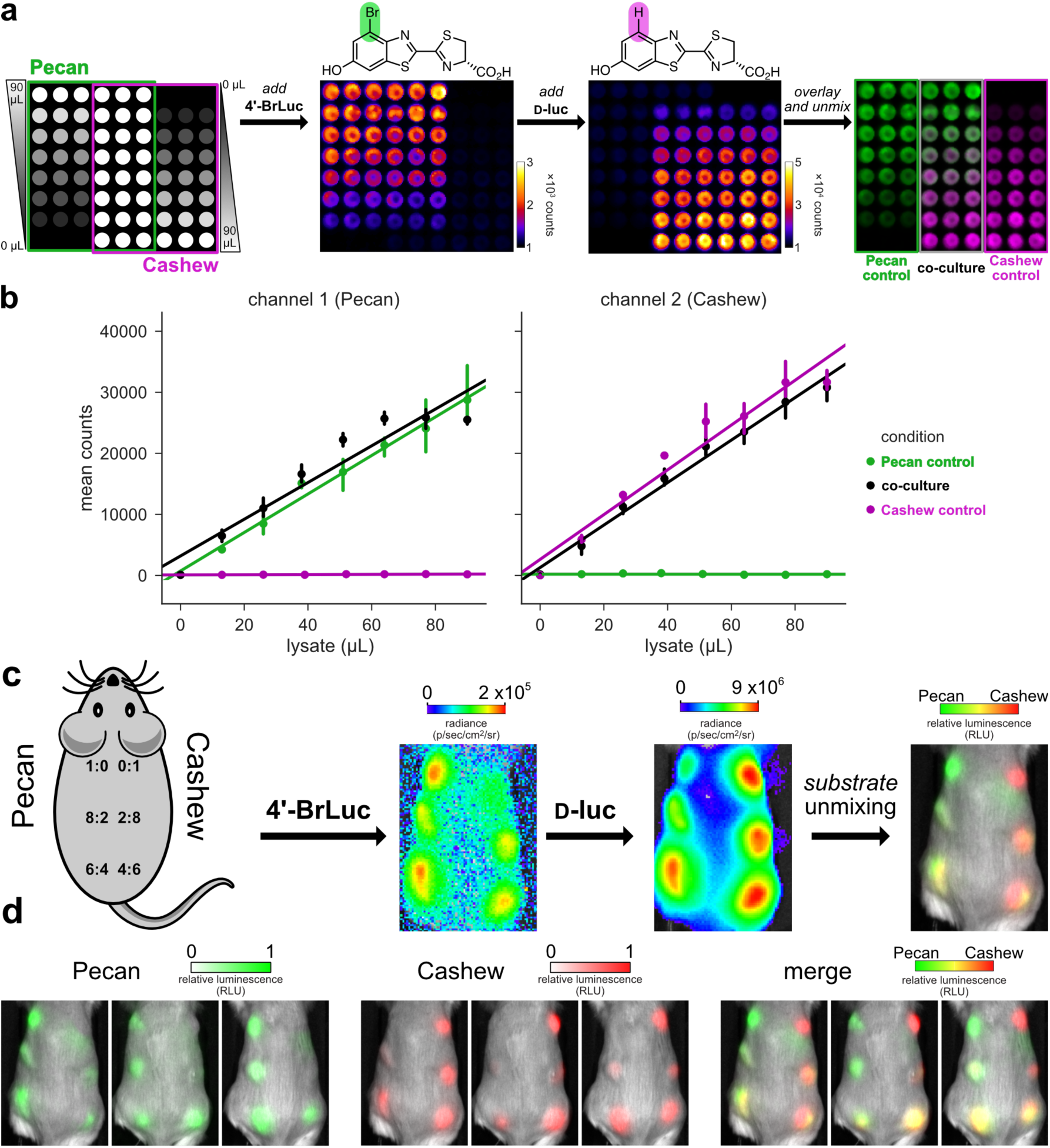
Rapid BLI *in vitro* and *in vivo*. (a) Pecan and Cashew were plated in a gradient fashion (as shown). The samples were treated with 4’-BrLuc (100 μM), followed by D-luc (100 μM). Raw images were acquired after each substrate addition. The substrate-specific signals were unmixed, assigned false colors and overlaid. (b) Quantification of the images from (a), fit via linear regression. Error bars represent the standard error of the mean for *n* = 3 experiments. (c-d) Orthogonal bioluminescent probes can be distinguished in mice. (c) Ratios of Cashew-and Pecan-expressing cells implanted in mice. Orthogonal substrates (65 mM) were administered sequentially via i.p. injection (100 μL). Images were acquired 35 min after each injection. (d) Unmixed channels for each mouse replicate are shown. Color bars indicate normalized luminescence values.

The rapid multicomponent BLI protocol was further applied *in vivo*. Pecan- and Cashew-expressing cells were mixed in varying ratios and implanted in mice (10^6^ cells per site, Fig 2c). Upon injection of 4’-BrLuc, Pecan-expressing cells were readily visualized (Fig 2d). Prior to substrate clearance, the brighter luciferin (D-luc) was injected. Signal was then observed from Cashew-expressing cells. Substrate unmixing revealed the expected distributions of Cashew- and Pecan-expressing cells (Figs 2d, S7). Notably, the two-component imaging session was complete in ∼1 hour, a significant improvement from the 6-24 h imaging window typically used with other substrate-resolved probes.^11^ The relative composition of the cell masses could be readily visualized. Like conventional BLI, the unmixing algorithm cannot provide *absolute* quantification of bioluminescent signals in animals. Rather, the *relative* amounts of signal are easily discerned and tracked. These measures are often paramount in optical imaging *in vivo* and similar intensiometric measures are routinely used in imaging with fluorescent sensors.^18-19^

Rapid BLI is generalizable to multiple luciferase reporters that use unique substrates and exhibit a range of intensities. To demonstrate with a set of three probes, we used Cashew and Pecan in combination with Antares, a recently reported marine luciferase variant.^20^ Cashew and Pecan derive from the insect luciferase family, and are thus immediately orthogonal to luciferases (like Antares) that use vastly distinct luciferins (in this case, furimazine).^6^ Antares also exhibits markedly faster substrate turnover than Cashew, rendering it brighter and intensity resolved from the other two reporters. We thus reasoned that the three orthogonal luciferases could be rapidly differentiated by first applying 4’-BrLuc, followed by D-luc, then furimazine to layer in signal from Pecan, Cashew, and Antares, respectively. When the cells were combined and imaged together, the three reporters could be rapidly visualized (<15 minutes) following sequential substrate addition (Fig 3). Triplet imaging was also readily achieved using other combinations of engineered and native luciferases (Fig S8). Historically, triple-component BLI has been difficult to execute due to a lack of streamlined protocols.^21^

**Figure 3.**
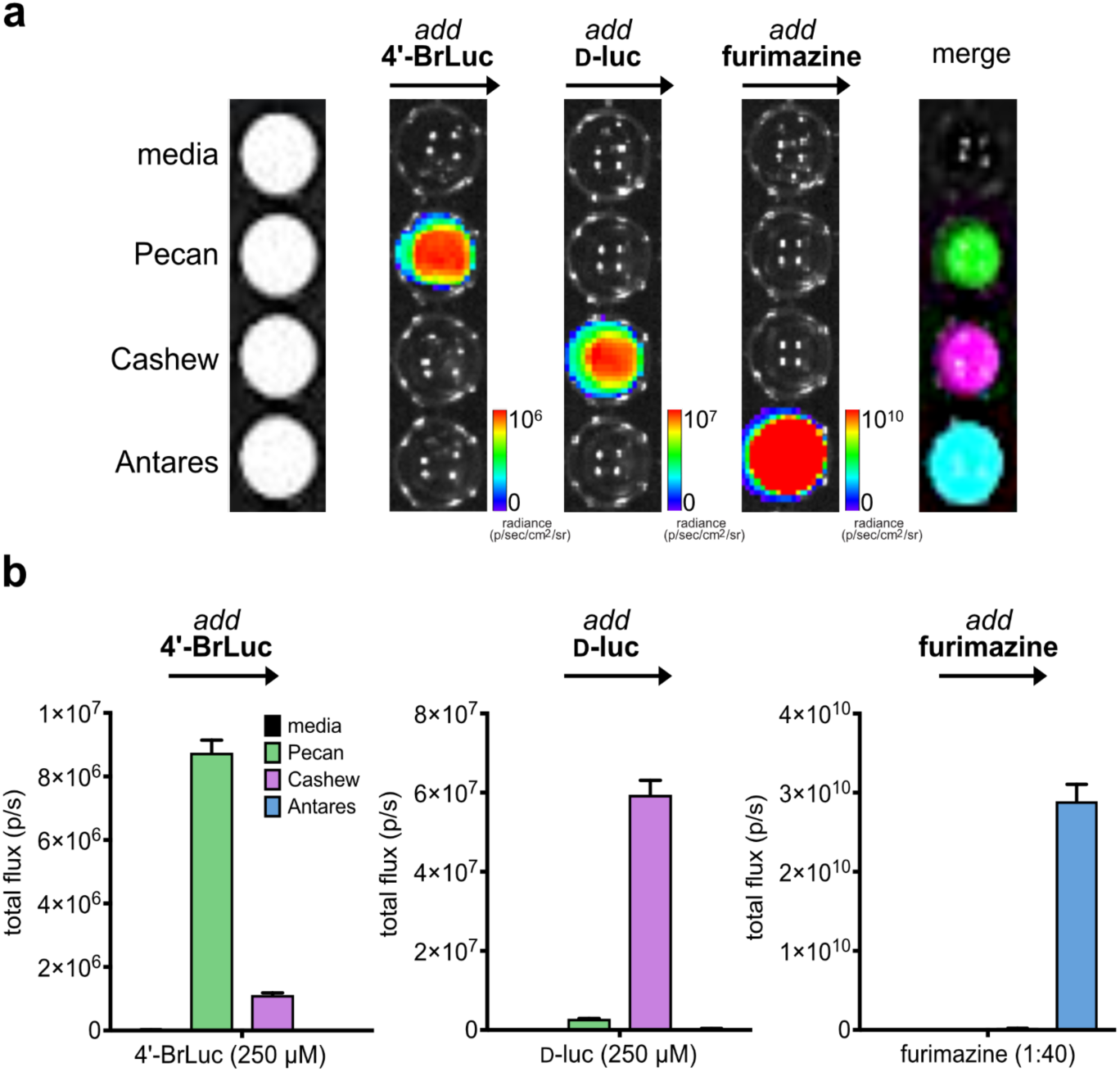
Rapid BLI with three luciferases and luciferins. (a) Cells expressing Pecan, Cashew, Antares, or no luciferase (control) were plated in a 96-well plate. Sequential substrate administration (4’-BrLuc, followed by D-luc, then furimazine) and unmixing enabled three-component imaging. Data are representative of *n* = 3 replicates. (b) Quantified photon outputs for the images in (a). Error bars represent the standard error of the mean for *n* = 3 experiments.

In conclusion, we developed a strategy for rapid multicomponent bioluminescence imaging based on substrate unmixing. This method takes advantage of both substrate and intensity resolution to resolve mixtures of reporters. We validated the approach in lysate, live cells, and mouse models. Probe differentiation is unaffected by tissue depth or location, parameters that have historically hindered efforts to resolve colors in large organisms. Like conventional BLI, the unmixing algorithm enables relative amounts of signal to be tracked over time in physiologically relevant environments. We anticipate that this approach will enable a range of applications, including monitoring multiple cell types and gene expression profiles. The development of additional intensiometric probes will further increase the number of bioluminescent probes that can be rapidly imaged in tandem. An expanded toolkit for BLI will enable longstanding questions regarding multicellular interactions to be addressed.

## ACKNOWLEDGEMENTS

This work was supported by the U.S. National Institutes of Health (R01 GM107630 to J.A.P.). W.B.P. was supported by the National Science Foundation via the BEST IGERT program (DGE-1144901) and an Allergan Graduate Fellowship. K.A.J. was supported by an institutional Chemical and Structural Biology Training Grant predoctoral fellowship (T32-GM10856). Z.Y. and C.M.R. were supported by National Science Foundation Graduate Research Fellowships. We thank members of the Prescher laboratory for helpful discussions.

## MATERIALS AND METHODS

### Reagents

All reagents purchased from commercial suppliers were of analytical grade and used without further purification. 4’-BrLuc, 4’-MorphoLuc, 7’-MorPipLuc, and 7’-DMAMeLuc were prepared and used as previously described.^10-11^

### General bioluminescence imaging

Assays were performed in black 96-well plates (Greiner Bio One). Plates were imaged in a light-proof chamber (IVIS Lumina, Xenogen) equipped with a CCD camera (chilled to –90 °C). The stage was kept at 37 °C during imaging experiments, and the camera was controlled using standard Living Image software. Exposure times ranged from 1 s to 5 min, and data binning levels were set to small or medium. Post-acquisition, regions of interest were selected for quantification. Total flux and radiance values were analyzed using Living Image software or ImageJ (NIH).

### Bacterial lysate analysis of luciferase mutants

Bacterial cell stocks (stored in glycerol) containing the mutations of interest (in pET vectors) were streaked on agar plates containing kanamycin. After overnight growth, colonies were picked and expanded overnight. Portions of the cultures (100 µL) were added to 5 mL of LB (kan) and luciferase expression was induced as described previously in Jones, *et al*.^10^

### Substrate unmixing

Substrate unmixing experiments were designed such that a “positive” sample for each enzyme was present in the image to be acquired. For *in vitro* experiments, “positive” wells comprised pure enzyme. In mouse experiments, “positive” cells were also included as a reference. Images were acquired as a series following each substrate addition. Thus, an image was generated for each enzyme/substrate pair. Linear unmixing was conducted using ImageJ (installed under the FIJI package). Luminescence images containing the raw CCD counts (as TIFF files) were loaded into FIJI and subjected to a 2-pixel median filter to remove any cosmic noise. Next, the signal at each pixel was min-max scaled to lie between 0 and 65535 (the maximum value that can be stored in a 16-bit image). As a result, the brightest pixel in each image had a value of 65535, and the dimmest had a value of 0. Images were then stacked, and an additional image containing the maximum value of each of the stacked images was computed (as a Z projection). This new image was added to the stack, and signals were unmixed using the ImageJ plugin developed by Gammon, *et al*.^14^ In the plugin, regions of interest (ROIs) for each luciferase were drawn around the “pure” areas of the image described above. Each ROI was drawn individually and added to the list by clicking “add.” Once all enzymes were added, “Unmix” was used to unmix the images. Pseudocolors were assigned in FIJI through the “Merge Channels” tool.

### Mammalian plasmid construction

Pecan- and Cashew-expressing DB7 cells were prepared via CRISPR gene insertion. The relevant luciferase genes (luciferase-G4SX2-FP-T2A-Puro) were amplified and inserted into CRISPR AAVS1 donor plasmids (courtesy of Drs. Theresa Loveless and Chang Liu, UCI). Cashew and Pecan inserts were amplified from pET vectors using the following primers: 5’-TGGCTAGCGCTACCGGTCGCCACCTCTAGAATGGAAGACGCCAAAAACATAAAGAAAGG-3’ and 5’-GCGGAAAGATCGCCGTGGGCGGAGGCGGGTCTGGGGGCGGAGGCTCT-3’

Antares inserts were amplified with the following primers: 5’-GCTAGCGCTACCGGTCGCCACCTCTAGAATGCGGGGTTCTCATCATCATCATC-3’ and 5’-TGCCTCTGCCCTCGCCGCTGCCCTCGAGCTTGTACAGCTCGTCCATGCCTCCG-3’ Linearized vectors were generated via digestion with restriction enzymes *Xba*I and *Xho*I (New England BioLabs). The linearized vectors were combined with the appropriate luciferase insert by Gibson assembly. A portion of the reactions (3.0 µL) was directly transformed into XL1 competent *E. coli* cells. Sequencing analysis confirmed successful plasmid generation.

### Mammalian cell culture and imaging

DB7 cells (courtesy of the Contag laboratory, Stanford) were cultured in DMEM (Corning) supplemented with 10% (vol/vol) fetal bovine serum (FBS, Life Technologies), penicillin (100 U/mL), and streptomycin (100 µg/mL). Cells were maintained in a 5% CO_2_ water-saturated incubator at 37 °C. To create stable lines expressing mutant luciferases, DB7 cells were transfected with the AAVS1 mutant luciferase donor plasmid, Cas9 (addgene #41815), and AAVS1 sgRNA (addgene #53370) using lipofectamine. The mutant luciferases were integrated into the first locus of AAVS1 through homologous recombination. Transfected cells were then treated with puromycin (20 µg/mL) and FACS sorted at the Institute for Immunology Flow Cytometry Core (UCI).

DB7 cells stably expressing luciferases were added to black 96-well plates (1 x 10^5^ cells per well). Stock solutions of 4’-BrLuc and D-luc (5 mM in PBS) were added to each well (250-500 µM final concentration). A solution of furimazine (1:40-1:100 dilution of the commercial stock, Promega) was then added. Sequential imaging was performed as described in the General bioluminescence imaging section (above).

### *In vivo* imaging of orthogonal luciferase-luciferin pairs

Mouse experiments were approved by the UC Irvine Animal Care and Use Committee. FVB/NJ mice (The Jackson Laboratory) received subcutaneous dorsal injections of 1×10^6^ DB7 mutant luciferase expressing cells. After 24 h, animals were anesthetized (2% isoflurane) and placed on a warmed (37 °C) stage for imaging. To eliminate movement between injections, one leg of each animal was secured to the imaging platform with black tape. Each mouse received an i.p. injection of luciferin (65 mM, 100 µL per mouse) under the non-secured leg. Images were acquired with 5 min exposure times for 35 min using the General bioluminescence procedure. For sequential imaging, mice were immediately injected with the second substrate and imaged for an additional 35 min. Bioluminescent output was quantified as above.

## SUPPORTING INFORMATION

**Figure S1.**
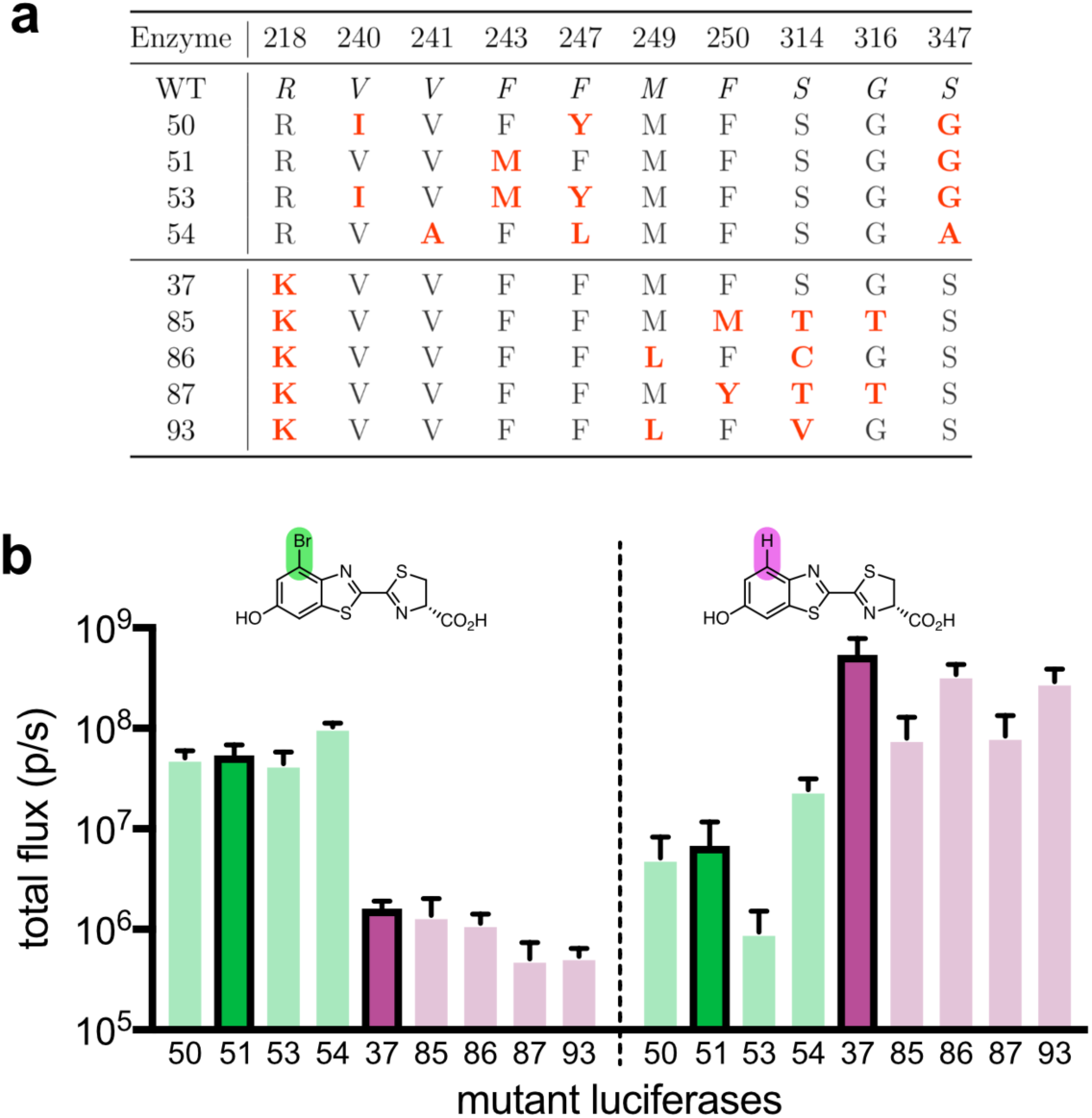
Identifying intensity resolved orthogonal pairs (a) Mutants considered for rapid BLI with 4’-BrLuc and D-luc. Mutants 50, 51, 53, and 54 prefer 4’-BrLuc, while 37, 85, 86, 87, and 93 prefer D-luc.^11^ (b) Candidate luciferases were expressed in bacteria and screened with 100 µM 4’-BrLuc or D-luc. Mutants exhibited orthogonal substrate use, with >10-fold substrate preference observed in most cases. The “winning combination” – mutants 51 and 37 – were also intensity resolved. Error bars represent the standard error of the mean for *n* = 3 experiments.

**Figure S2.**
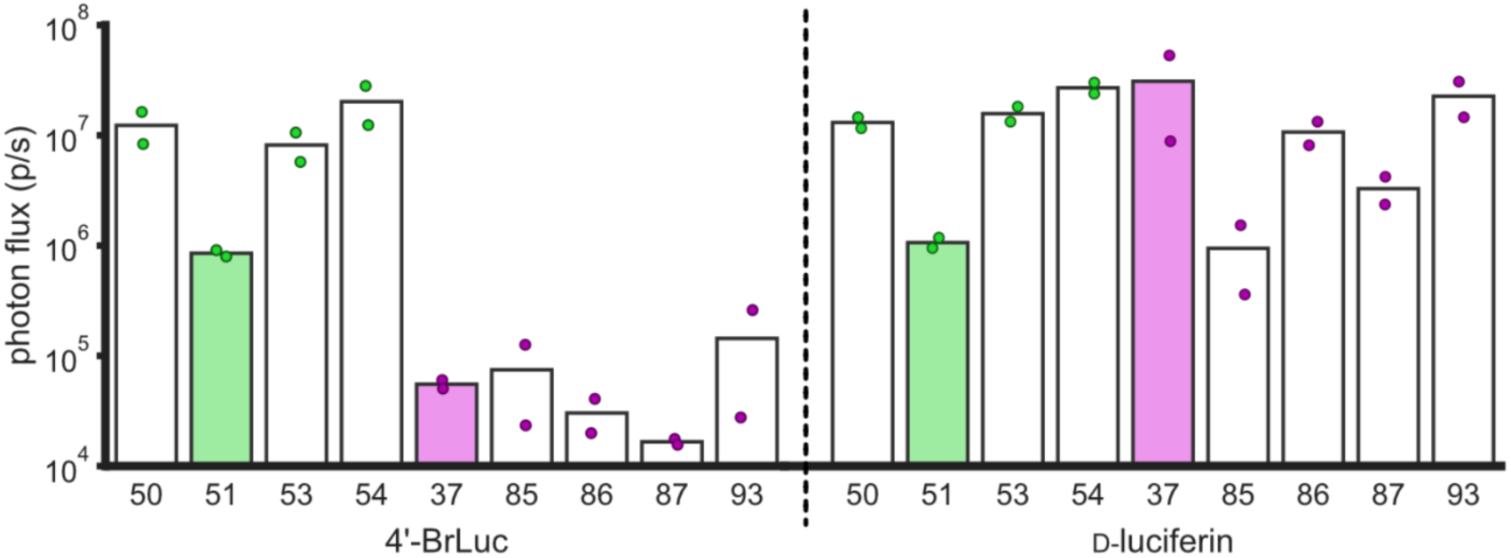
Verifying orthogonality and substrate resolution *in vivo*. DB7 cells expressing different mutant luciferases were implanted into the backs of mice. Sequential application of 4’-BrLuc and D-luc enabled identification of optimal mutant luciferase combinations for multicomponent imaging. Photon flux values from images were quantified and plotted. Mutant luciferases 51 and 37 were identified as optimal candidates for rapid multicomponent BLI.

**Figure S3.**
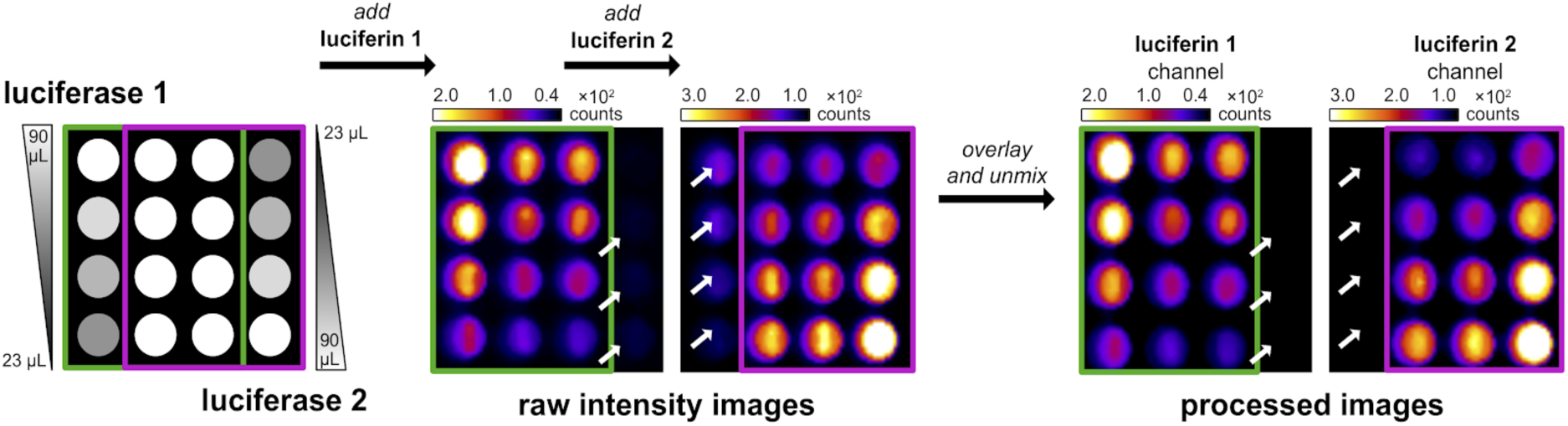
Residual signals are removed by substrate unmixing. Gradients of mutant luciferases in bacterial lysate were plated in a 4×4 matrix. The corresponding luciferins were added sequentially. Pixels containing residual signal are highlighted by the white arrows. These signals are removed upon imaging processing.

**Figure S4.**
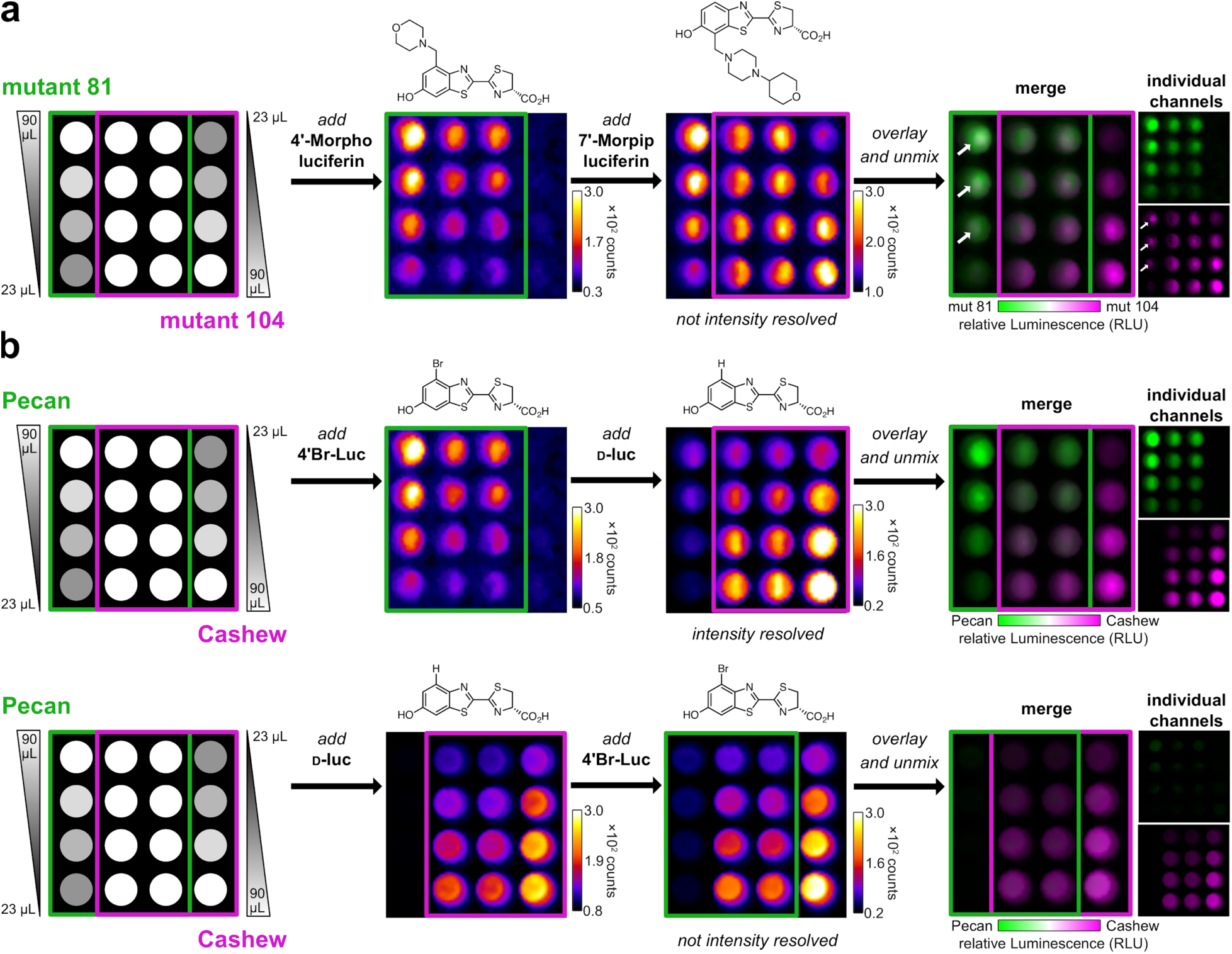
Substrate unmixing requires probes that are intensity resolved. (a) An orthogonal pair comprising mutant 81/4’-Morpho luciferin and mutant 104/7’-Morpip luciferin are not intensity resolved, and thus not amenable to rapid BLI. Gradients of the mutants (expressed in bacterial lysate) were plated in a 4×4 matrix. 4’-Morpho luciferin (the preferred substrate for mutant 81) was then administered, followed by 7’-Morpip luciferin (the preferred substrate for mutant 104). Substrate unmixing was not successful. Strong residual signal (from 4’-Mopho luciferin) in the the 7’-Morpip luciferin image can be observed. The presence of white pixels in the merged image (arrows) is not consistent with the composition of the well (only one luciferase was present). (b) Gradients of Cashew and Pecan in bacterial lysate were plated in a 4×4 square. When the dimmer analog (4’-BrLuc) was added prior to the brighter one (D-luc), the signals can be readily unmixed (top). When D-luc was added first, though, the signals cannot be distinguished (bottom).

**Figure S5.**
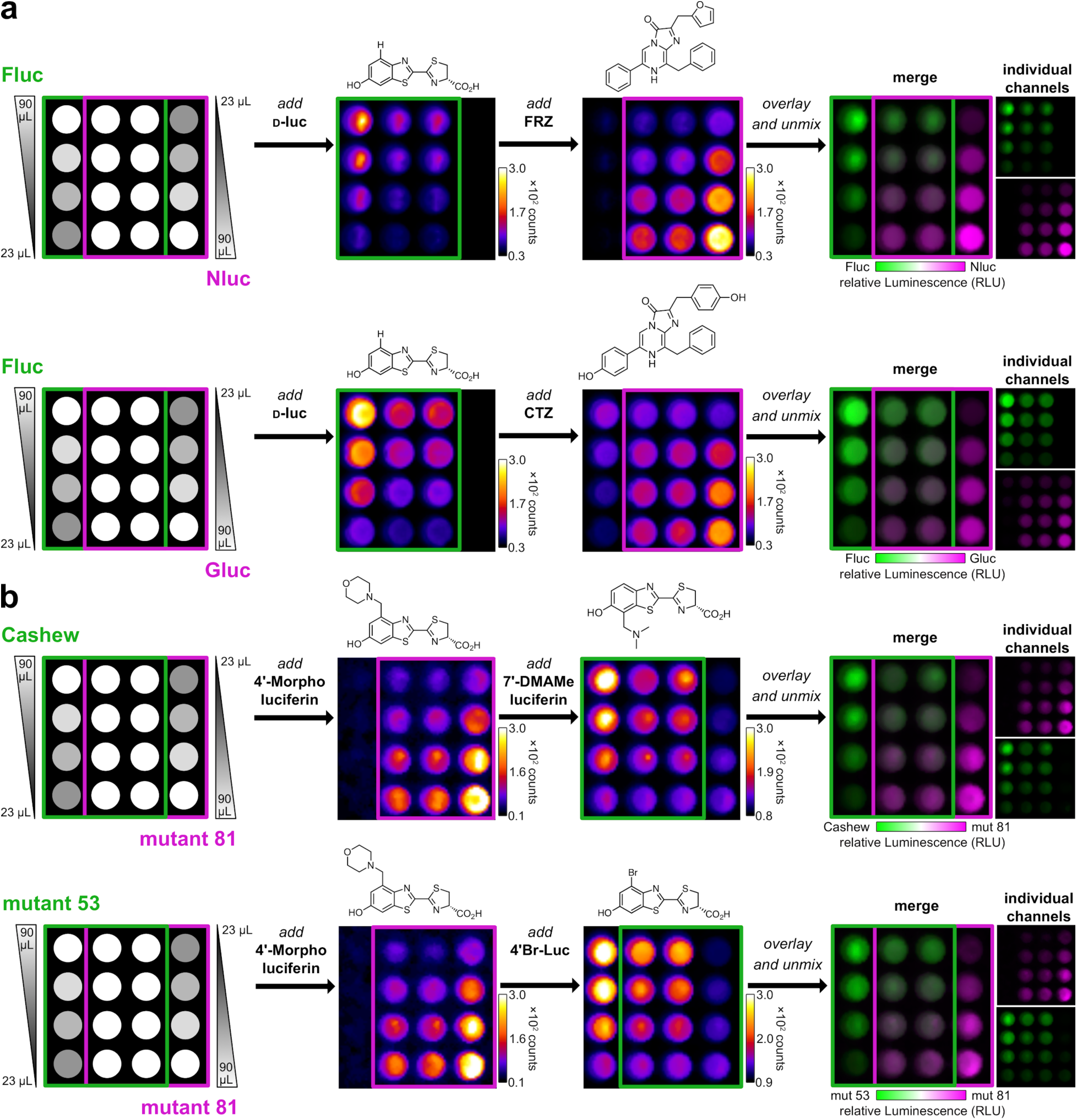
Multiple orthogonal pairs can be rapidly unmixed. Gradients of (a) native luciferases or (b) engineered variants were plated as shown. The corresponding substrates were administered, beginning with the dimmest luciferin. Images were acquired after each addition. The raw data were stacked and unmixed. Established reporters examined include firefly luciferase (Fluc), NanoLuciferase (Nluc), and *Gaussia* luciferase (Gluc). The corresponding luciferins for each reporter are shown. The engineered luciferase-luciferin pairs include mutant 81 (and its corresponding substrate 4’-MorphoLuc), Cashew, and mutant 53 (both of which can process 4’-BrLuc).

**Figure S6.**
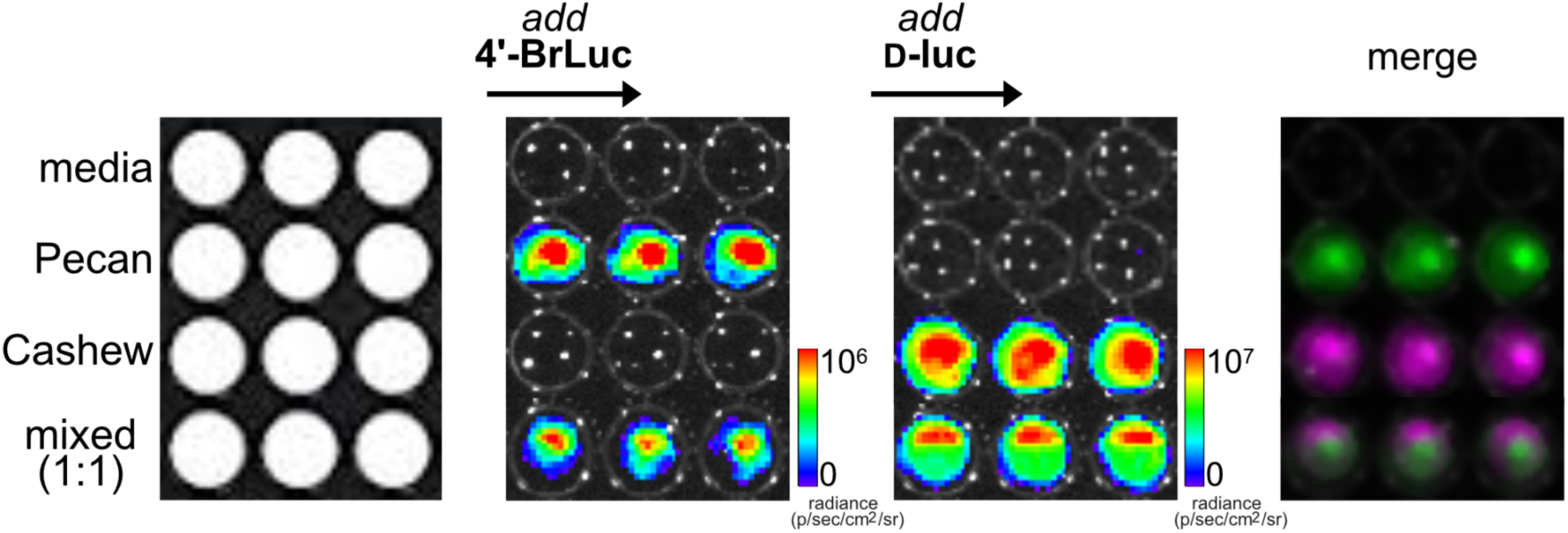
Sequential substrate administration enables multicomponent bioluminescence imaging *in cellulo.* DB7 cells expressing Cashew, Pecan, or no luciferase (media) were plated (1 × 10^5^ cells/well). Some wells contained a 1:1 mixture Cashew- or Pecan-expressing cells (5 × 10^4^ of each cell type per well). All samples were first treated with 4’-BrLuc, followed by D-luc. Images were acquired after each substrate addition. Raw photon values are shown, along with the merged image following substate unmixing.

**Figure S7.**
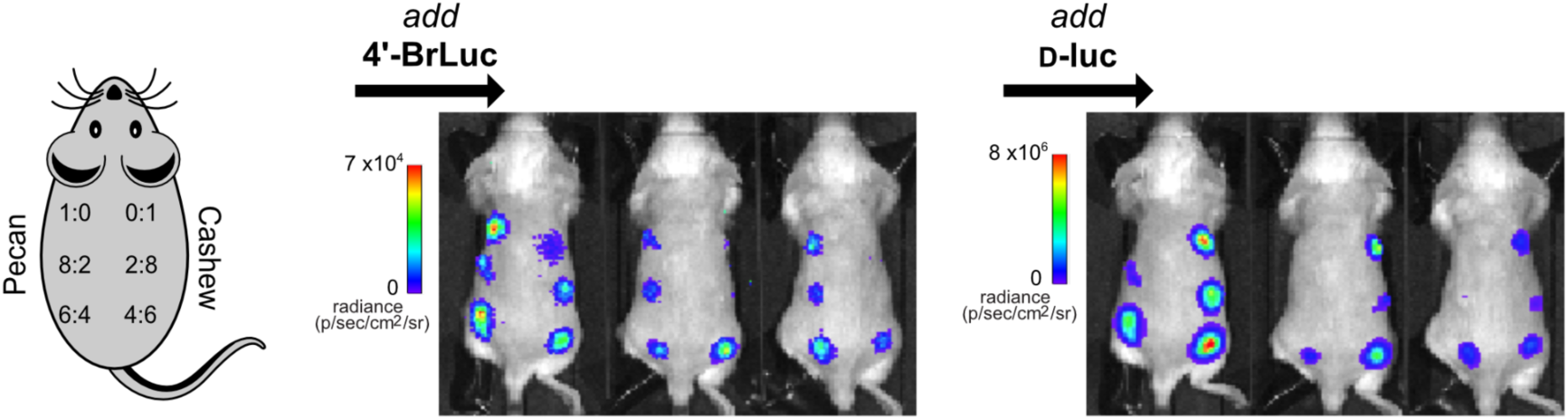
Multicomponent BLI in mouse models. Images used to generate the false colored pictures in Fig 2d are shown. Sequential application of 4’-BrLuc and D-luc enabled different ratios of Pecan- and Cashew-expressing cells to be visualized.

**Figure S8.**
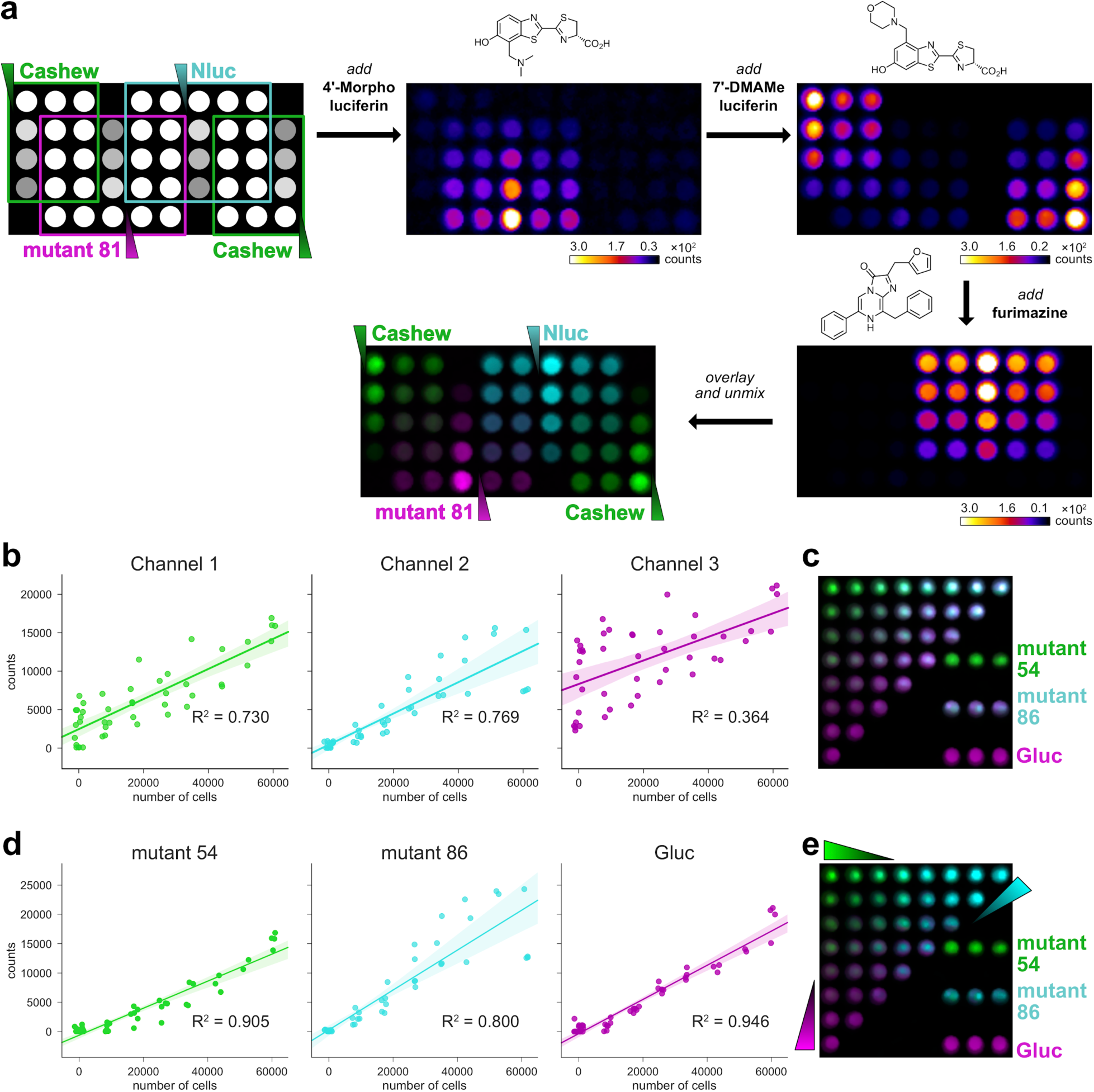
Three orthogonal probes can be distinguished in bacterial lysate and mammalian cells. (a) Gradients of luciferases in bacterial lysate were plated in a 96-well plate. 7’-DMAMeLuc luciferin (250 µM), 4’-Morpho luciferin (250 µM), furimazine (1:100 dilution of commercial stock) were added in sequence. Images were acquired after each addition, and the raw data were stacked and unmixed. (b-e) Gradients of cells expressing luciferase mutants 51 and 86, or *Gaussia* luciferase were plated in a triangle, with 60,000 cells per well. 4’-BrLuc (500 µM), D-luc (500 µM), and coelenterazine (40 µM) were added in sequence. (b) Quantification of each channel from (c) fit via linear regression. The shaded area represents the 95% confidence interval of the fit. (c) Overlay of raw signal from mixed images. (d) Quantification of each channel from the unmixed image in (e) fit via linear regression. The shaded area represents the 95% confidence interval of the fit. (e) Overlay of the unmixed channels.

## Spectral Unmixing Algorithm

### Linear Unmixing

Spectral unmixing was developed to automate the analysis of images that contain multiple components of overlapping spectra. It is commonly used in fluorescence imaging to deconvolute fluorophores that are not spectrally resolved.^1^ Due to resolution constraints, each pixel in a fluorescence image has the chance to contain different fluorophores. A fundamental assumption of linear unmixing is that the signal of each pixel in an image is a linear combination of the absolute contents of that pixel.

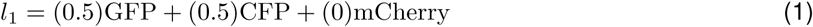

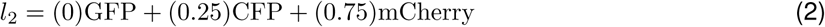

Where *l*_1_ and *l*_2_ represent the fractional composition of individual pixels in an image. These pixels can be represented as vectors where each component of the vector is a different fluorophore.

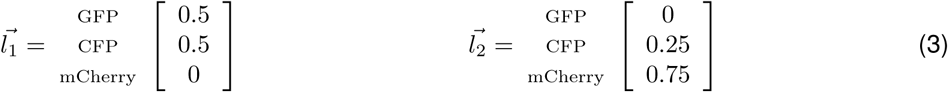

Each pixel in these fluorescence images is collected with spectral information (with filters, for example). We can represent the raw pixel data as a different vector 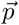 sities, *i*, at each wavelength (or wavelength range) that was measured:

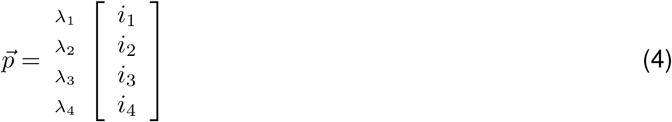

Where *λ*_1_ to *λ*_4_ are measurements at each wavelength, and *i*_1_ to *i*_4_ are intensities measured at each of those wavelengths. Each component of this vector is dependent on the fraction of fluorescent labels that comprise the pixel, 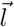, and the spectrum of each of those fluorophores, *K*. Thus, for each pixel in an image:

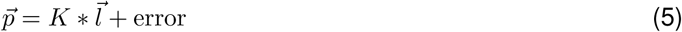

Written out, this would look like:

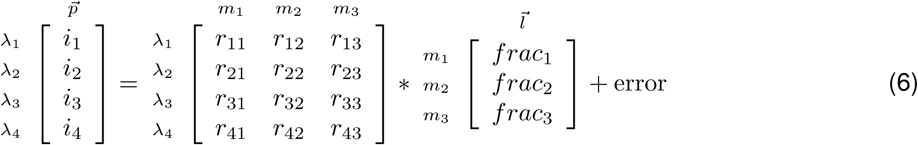

Where *m*_1_ to *m*_3_ are the spectra of the various fluorescent labels that might comprise the pixel, for example, CFP, GFP, and mCherry, and *frac*_1_, *frac*_2_, and *frac*_3_ are the fractional amounts of each of these fluorophores (as in 3). Thus, solving for 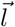:

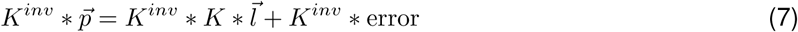

and rearranging:

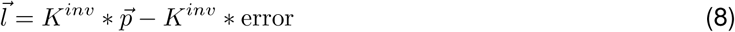

For an entire image (list of pixels), 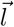 and 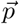 become matrices *L* and *P*:

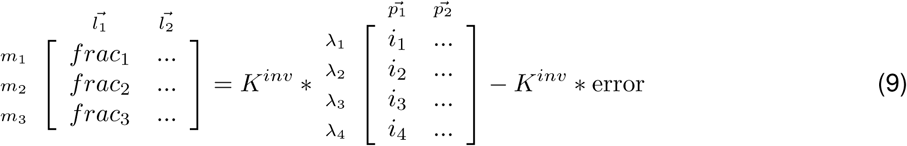

or

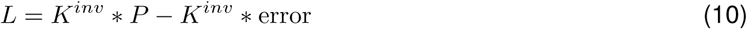

By arranging the equation in this way, we can measure the spectrum at each pixel, 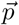 to calculate the fractional makeup of each pixel, 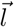.

### Bioluminescence Imaging (previous work)

Similarly, these concepts can be translated to bioluminescence imaging (BLI). In the case of BLI, the signal from each pixel represents a mixture of the various labeled cell types. Previously, linear unmixing with bioluminescence has been used in a similar fashion to fluorescence imaging; the various components of 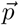 were wavelengths (or ranges of wavelengths) measured with filters. ^2–4^

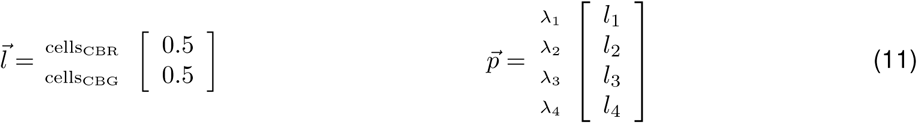

Where cells_CBR_ and cells_CBG_ are cells expressing click beetle red luciferase and click beetle green luciferase respectively, and *λ*_1_ to *λ*_4_ are various filters used by the *in vivo* imaging system.

### Substrate-resolved Bioluminescence (this work)

This work utilizes the same basic algorithm (equation 8), and (in a similar fashion as equation 11) defines the components of 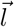 as cells labeled with various luciferase mutants. The major difference from all other applications of linear unmixing is that the wavelengths of 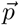 have been replaced with various luciferin substrates. Thus, instead of taking images at a variety of wavelengths, we are imaging after the addition of each luciferin substrate.

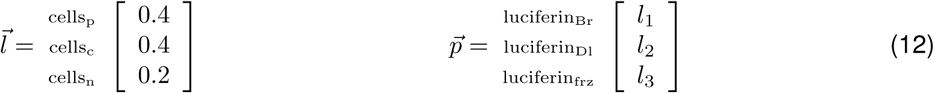

Where cells_p_, cells_c_, and cells_n_ are cells expressing pecan, cashew, and nanoluc respectively, and luciferin_Br_, luciferin_Dl_, and luciferin_frz_ are the substrates 4’-BrLuc, D-Luc, and furimazine respectively. Thus, we can rewrite equation 9 as:

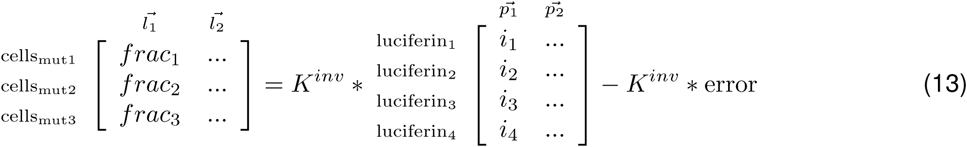

In this implementation, *K* can be determined by measuring the response of each mutant individually across the sequential addition of luciferin substrates:

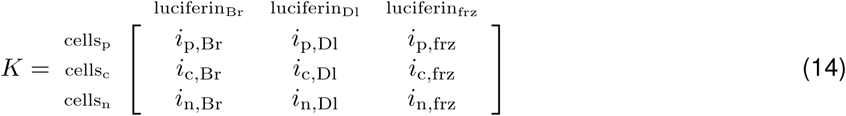

In practice, finding *K* is a matter of including calibration sites in the image where each mutant is segregated, and receives each compound in the same manner as the rest of the experiment (just as if filters were being applied to calibration wells to determine spectral response).

## Intensity Resolution

The technique described herein for multicomponent bioluminescence imaging via sequential substrate administration also relies on *intensity* resolution amongst the substrates. In fact, we have found that for the success of the technique, intensity resolution is just as important as substrate resolution. We define intensity resolution as the difference in brightness between two probes. Probes that are intensity resolved enable sequential administration in a single imaging session because the highest possible signal in the probe that is administered first minimally overlaps with the lowest possible signal in the second probe that is administered. If the three probes, *p*, are administered in sequence (1–3), the total signal intensity, *I*, from the contributions of each individual probe, *i*, following each addition can be illustrated as follows:

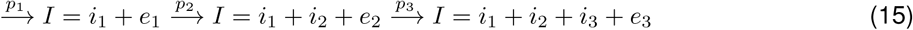

Where *e* is the random error associated with each image acquired. In order to be able to resolve all the individual signals, it is necessary that:

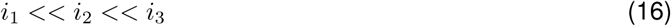

When the intensity resolutions (differences) between each probe are large enough that the signal of the previous probe is similar to the error, the signals can be resolved. Written out:

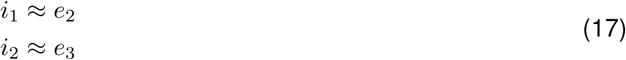

This enables us to eliminate the residual intensity terms from equation 15 to give:

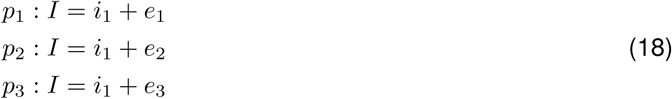

Any systematic error that is contributed by the previous probe can be eliminated through linear unmixing, described above. Large amounts of intensity overlaps cannot be solved by unmixing, however. If the relationship in the above equation (16) is not true, we risk not having enough information to resolve the individual signals.

